# Spatial and Temporal Analysis of SARS-CoV-2 Genome Evolutionary Patterns

**DOI:** 10.1101/2023.06.21.545910

**Authors:** Shubhangi Gupta, Deepanshu Gupta, Sonika Bhatnagar

## Abstract

The spread of SARS-CoV-2 virus accompanied by availability of abundant sequence data publicly, provides a window for determining the spatio-temporal patterns of viral evolution in response to vaccination. In this study, SARS-CoV-2 genome sequences were collected from seven countries in the period January 2020-December 2022. The sequences were classified into three phases, namely: pre-vaccination, post-vaccination, and recent period. Comparison was performed between these phases based on parameters like mutation rates, selection pressure (dN/dS ratio), and transition to transversion ratios (Ti/Tv). Similar comparisons were performed among SARS-CoV-2 variants. Statistical significance was tested using Graphpad unpaired t-test. The comparative analysis showed an increase in the percent genomic mutation rates post-vaccination and in recent periods across different countries from the pre-vaccination phase. The dN/dS ratios showed positive selection that increased after vaccination, and the Ti/Tv ratios decreased after vaccination. C→U and G→U were the most frequent transitions and transversions in all the countries. However, U→G was the most frequent transversion in recent period. The Omicron variant had the highest genomic mutation rates, while Delta showed the highest dN/dS ratio. Mutation rates were highest in NSP3, S, N and NSP12b before and increased further after vaccination. NSP4 showed the largest change in mutation rates after vaccination. N, ORF8, ORF3a and ORF10 were under highest positive selection before vaccination. They were overtaken by E, S and NSP1 in the after vaccination as well as recent sequences, with the largest change observed in NSP1. Protein-wise dN/dS ratio was also seen to vary across the different variants.

**Importance:** Irrespective of the different vaccine technologies used, geographical regions and host genetics, variations in the SARS-CoV-2 genome have maintained similar patterns worldwide. To the best of our knowledge, there exists no other large-scale study of the genomic and protein-wise mutation patterns during the time course of evolution in different countries. Analysing the SARS-CoV-2 evolution patterns in response to spatial, temporal, and biological signals is important for diagnostics, therapeutics, and pharmacovigilance of SARS-CoV-2.

## INTRODUCTION

Coronavirus disease 2019 (COVID-19) emerged as an epidemic with its initial spread in Wuhan (China) and was declared as a pandemic in March 2020 by the World Health Organization. Over 650 million confirmed cases of COVID-19, with more than 6 million deaths have been reported worldwide by WHO as of 15 May 2022 (1). In India, the first case of COVID-19 was reported in January 2020. An exponential increase in the number of cases from March 2020 to September 2020 marked the first wave. The second wave was marked by a sudden increase in COVID-19 cases from March 2021 and high death rates (2). The Omicron variant of the causative SARS-CoV-2 characterized the third wave of COVID-19 with higher transmissibility but lower death rates (3).

SARS-CoV-2 has continuously evolved over time, resulting in numerous genetic variants across the globe (4). These variants are classified into three main categories, i.e., Variant under monitoring (VUM), Variant of Interest (VOI), and Variant of concern (VOC) (5). The VUM includes the GH/G (S gene D614G variant) progeny.; VOI includes the Mu (lineage B.1.621) and Lambda (C.37); whereas VOC includes the alpha (lineage B.1.1.7), beta (lineage B.1.351), gamma (lineage P.1), delta (lineage B.1.617.2), and omicron variants (lineages BA1, BA2, BA4, and BA5) (5, 6).

The S protein is responsible for SARS-CoV-2 attachment and entry by binding to the ACE2 (7–10). Continuous mutations in the Spike increase virus adaptability for escaping vaccine treatment resulting in high survival rates and spread of the virus (11–14). Multiple mutations in the omicron variant S protein may also strengthen its interaction with ACE2, thereby leading to antibody escape (15, 16). Mutations in the S region of SARS-CoV-2 have led to epitope loss, resulting in escape from the vaccine treatment, the most frequent mutation being D614G (17, 18). D614G is also found in the S region of clades G/GR/GRY/GH/GV and has high human host infectivity rate due to efficient transmission (19, 20).

The SARS-CoV-2 genome contains 14 ORFs. Of these, *ORF1a* encodes the NSPs 1-11 while *ORF1b* encodes NSP12-16. Together NSPs 1-16 form the replicase-transcriptase complex. This is followed by 13 ORFs encoding the four main structural proteins, namely S, E, M, N and interspersed by 9 accessory factors (21). Various studies have been performed on genomic mutations in the SARS-CoV-2 virus. The ratio of dN/dS >1 indicates positive selection and has been reported in the S glycoprotein(22, 23). Comparative analysis of SARS-CoV, SARS-CoV-2 and MERS-CoV substitution models showed a positive evolution model along with higher dN/dS, due to the dominance of dN (24, 25). Studies of mutations in the diagnostic targets in COVID-19 have suggested that the *N* gene has the highest number of mutations (26, 27).

Analysis of 469 genome sequences from Indian patients led to the identification of 536 dN and dS mutations in the six genes; *ORF1ab*, *S*, *N*, *ORF3a*, *ORF7a*, and *ORF8* (28). A broad analysis showed 33 different mutations in 837 Indian SARS-CoV-2 whole genome sequence isolates, of which 18 were unique to India. S , N, NSP3, NSP12, and NSP2 coding genes showed novel mutations and dN was found to be more than dS by approximately 3 folds (29). Modelling of the epidemic with different strains and mutations showed the emergence of a virus with higher transmissibility and evolutionary adaptations (30).

In this study, we calculated the genomic rates of mutation and dN/dS ratios in SARS-CoV-2 genome sequences taken from seven countries and showed that they increased with time, whereas the Ti/Tv decreased. Similarly, these parameters were also estimated for different known SARS-CoV-2 variants, where Omicron variant sequences showed highest mutation rate as compared to the other variants, but delta showed the highest dN/dS ratio. The highest mutating protein along with their individual dN/dS ratios and mutation rates were determined within each country. NSP3, S, N and NSP12b had the highest genomic mutation rates both in before and after vaccination phase. NSP3 showed highest genomic mutation rates before vaccination, which was replaced by S in the after vaccination, while a significant rise was observed in the NSP4 genomic mutation rate. N, ORF8, ORF3a and ORF10 were under strong positive selection pressure before vaccination, whereas after vaccination, E, S and NSP1 showed strong positive selection pressure, with NSP1 showing a notable increase in the dN/dS ratio. The estimated properties showed similar patterns in genetic variability across all geographic regions, despite the use of different vaccine technologies. This implies that the forces of evolution have been uniform across multiple parameters. However, there is a definite change in the pattern of mutations before and after vaccination. While the highly mutating and positively selected genes are important for pharmacovigilance and vaccination, the negatively selected ones are important for diagnostics. This is the only study that has compared these different parameters in response to time, geographical region, and vaccination event.

## Methods

### Collection of genomic sequences of SARS-CoV-2

To perform a critical analysis of the data, genome sequences of SARS-CoV-2 were retrieved in FASTA format for different countries using the GISAID database (31, 32). The countries taken into consideration included India, England, Canada, Italy, France, USA (Washington) and the Netherlands.

Additionally, genome sequences of different SARS-CoV-2 variants like Alpha, Beta, Gamma, Lambda, Mu, Delta, and Omicron were also retrieved.

For the accuracy of genome sequences retrieved from the database, filters like complete sequence, high coverage, and complete collection date were applied. The sequences downloaded were high coverage, with < 1% Ns and < 0.05% non-synonymous singlets.

### Classifying the genome sequences retrieved

SARS-CoV-2 genome sequences submitted in the GISAID database from the seven countries were downloaded from three distinct periods. The genome sequences were classified into three categories as follows:

a. The sequences considered as pre-vaccination sequences had the sample collection date of at least one month before the start date of vaccination taken from the website of WHO and from the Government vaccination websites of the concerned countries. The datasets were taken in triplicates.
b. The start date of vaccination was different for each country considered. After the date of start of vaccination, a buffer period is required for the vaccine to reach the population. Therefore, sequences collected 5-6 months after the date of start of vaccination were considered as after vaccination sequences. The datasets were taken in triplicates.
c. A set of sequences that constitute a recent dataset were also taken. As the available number of recent sequences were less in number for many countries, triplicate datasets for this period could not be taken.
d. Additionally, approximately 1000 sequences of each SARS-CoV-2 variant were obtained using the “Variant” filter of GISAID.

### Sequence alignment and mutational data analysis

The first reported sequence, Wuhan-Hu-1 (Accession ID: NC045512), was taken as the reference for pre-vaccination, post-vaccination, and recent sequences. Thus, Wuhan-Hu-1 sequence was added as a reference for the alignment. The mutation rates of all the genomic sequences were calculated using COVID-19 genome annotator, an online web-based tool that aligns the input sequences in “nucmer” alignment tool and processes the alignment output using UNIX along with R scripts(33). The aligned sequences were compared with the reference genome (Wuhan-Hu-1) and amongst each other to find information about the mutational sites. The Jupyter notebook (34) was used in association with the “pandas” python library (35) to visualize and analyse the data obtained. The following values were calculated:

a. Percentage mutation rate The mutation rate has been calculated using the following equation: -

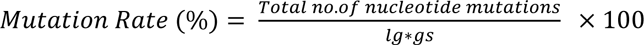

where ‘lg’ refers to the length of the data set taken, ‘gs’ refers to the length of the genome sequence and the total no. of mutations estimated with respect to number of single nucleotide mutations (36). Mutation rate is the measure that refers to the frequency of mutations per generation in the population or in an organism (37).
b. Selection pressure has been calculated as the dN/dS ratio. If dN/dS ratio exceeds unity (dN/dS > 1), the mutations are said to be occurring under positive selection which promotes the accumulation of beneficial mutations, whereas if dN/dS ratio is below unity (dN/dS < 1), the mutations are said to be occurring under negative selection that promotes mutations that are favouring selective removal of deleterious alleles (38, 39).
c. The Ti/Tv in the RNA virus has been calculated as no. of Ti (C↔U and A↔G) to the no. of Tv (A↔C, A↔U, G↔C and G↔U), determined for the pre-, post-, and recent vaccination sequence groups using the data obtained from the COVID-19 genome annotator.

### Statistical analysis

The statistical significance of the differences between the means was carried out. The unpaired t-test was performed on the % mutation rates, selection pressure and Ti/Tv ratios using GraphPad QuickCalcs (https://www.graphpad.com/quickcalcs/ttest1.cfm).

## Results and Discussion

In this work, we have carried out comparison of the SARS-CoV-2 genome sequences from seven different demographic regions, namely India, France, England, Canada, Italy, Netherlands, and USA (Washington D.C.). A total of 74,870 retrieved sequences were classified into three different time periods:

a. <u>Pre-vaccination phase</u>: Broadly, the pre-vaccination sequences taken were ranging from January 2020-December 2020 (GISAID Identifier: EPI_SET_230517nt).
b. <u>Post-vaccination phase</u>: The post-vaccination sequences taken were ranging from May 2021-April 2022 (GISAID Identifier: EPI_SET_230517va).
c. <u>Recent period</u>: The recent period sequences were taken from June 2022-December 2022 (GISAID Identifier: EPI_SET_230518oq).
d. Additionally, a total of 7209 sequences of different variants were also analysed. As new variants emerged sequentially over time, collection dates of the sequences considered for the SARS-CoV-2 variants ranged from November 2020 to January 2022 (GISAID Identifier: EPI_SET_230518xv).

The sequences were compared for % mutation rates, dN/dS ratio and Ti/Tv ratio. The analysis was carried out for the whole genome as well for every individual gene.

### SARS-CoV-2 genomic mutation rates increased over time

The % genomic mutation rates estimated for each country in the pre-vaccination, post-vaccination and the recent period are listed in Table 1. The genome mutation rates were calculated as (0.04 ± 0.02) % in the pre-vaccination phase. The average mutation rates increased to (0.17 ± 0.05) % in the post-vaccination period. A comparable increase was seen in all the seven countries studied. The mutation rates for sequences from a more recent interval were higher than the post-vaccination period. The mutation rate was found to be (0.28 ± 0.01) %, consistently across all the seven countries. Thus, there was an average increase of 3-4 folds in the % genomic mutation rates in all the countries after vaccination and 6-7 folds increase in the recent period in comparison with mutation rates in the pre-vaccination sequences.

**Table 1:**
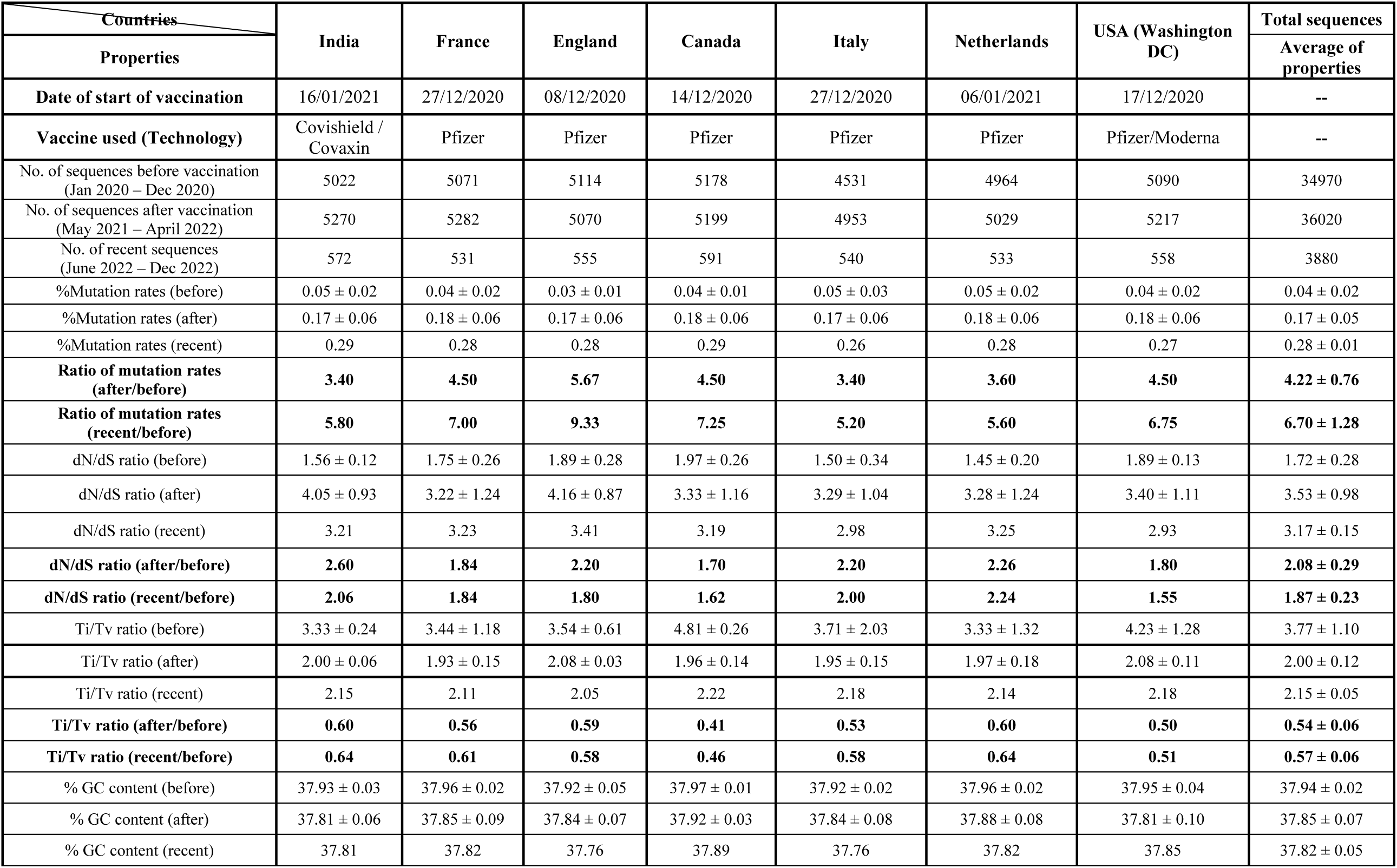
Data comparison of different countries before and after vaccination along with the recent data

Figure 1(A) shows the comparison of percent mutation rates within the countries in pre-vaccination, post-vaccination, and recent period. There was a similar increase in the % genomic mutation rates in the post-vaccination period in comparison with the pre-vaccination period across all the seven countries studied. This has increased further in the recent sequences. Despite the large variation in mutation rates, there was a definite increase in the post-vaccination sequences especially in the post-vaccination period. In the recent sequences, the mutation rates show a still higher trend in all the seven countries studied. The significance of the difference between the means was verified using unpaired t-test (P-value 0.0001; confidence interval of 95%) and is shown in Table 2. The differences in the percent genomic mutation rates were found to be extremely statistically significant between the three phases with a two-tailed p-value of less than 0.0001 (Table 2).

**Figure 1:**
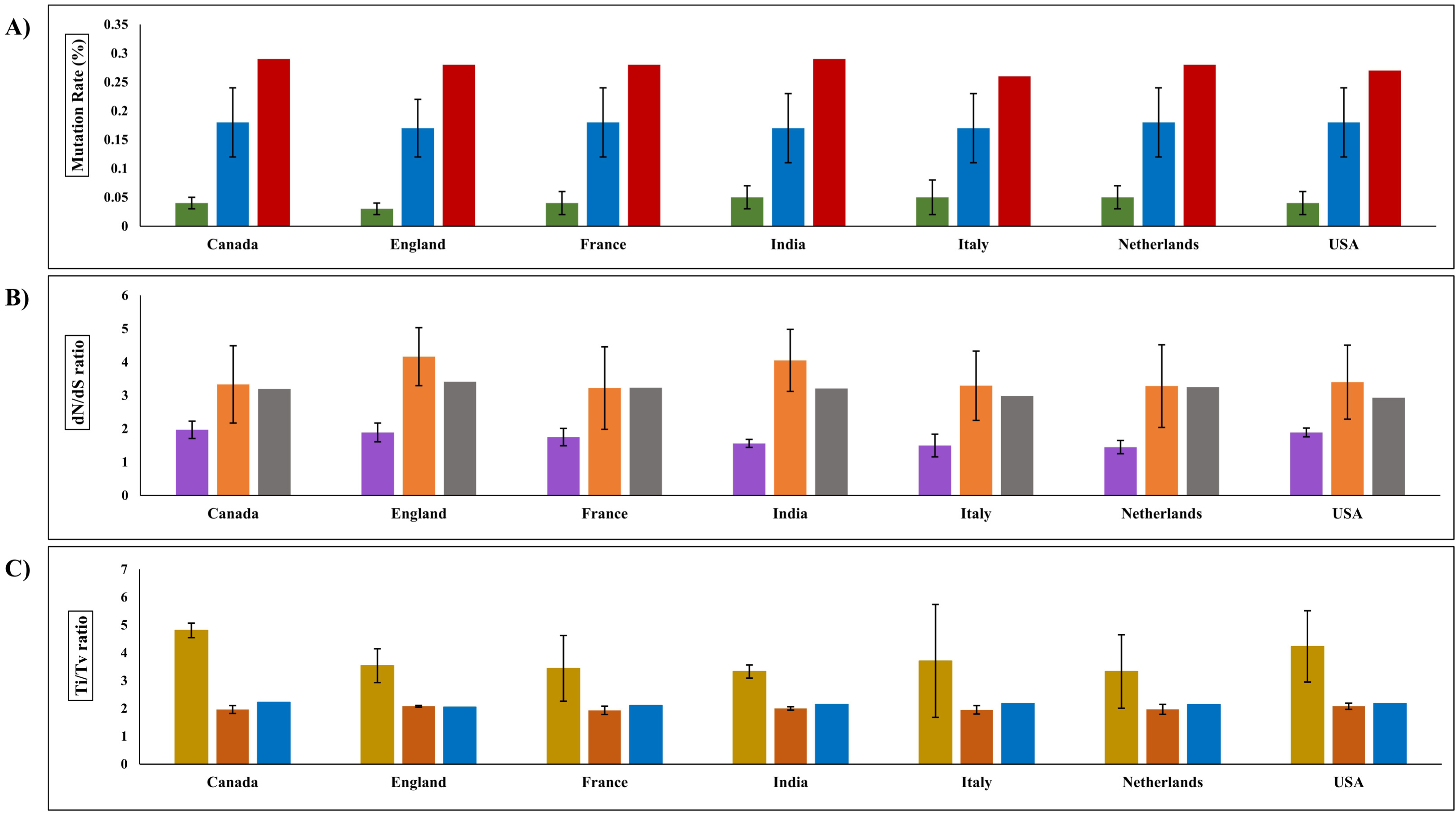
Comparison of mutation rates, dN/dS and Ti/Tv within the countries in the three phases. **A)** Mutation rates with respect to before (green), after vaccination (blue) and recent period (red); **B)** Selection pressure (dN/dS ratio) with respect to before (violet), after vaccination (orange) and recent period (grey); **C)** Transition to transversion (Ti/Tv) ratio with respect to before (yellow), after vaccination (pink) and recent period (brown). Black margins in the graph represent the standard deviation in the before and after vaccination.

**Table 2:**
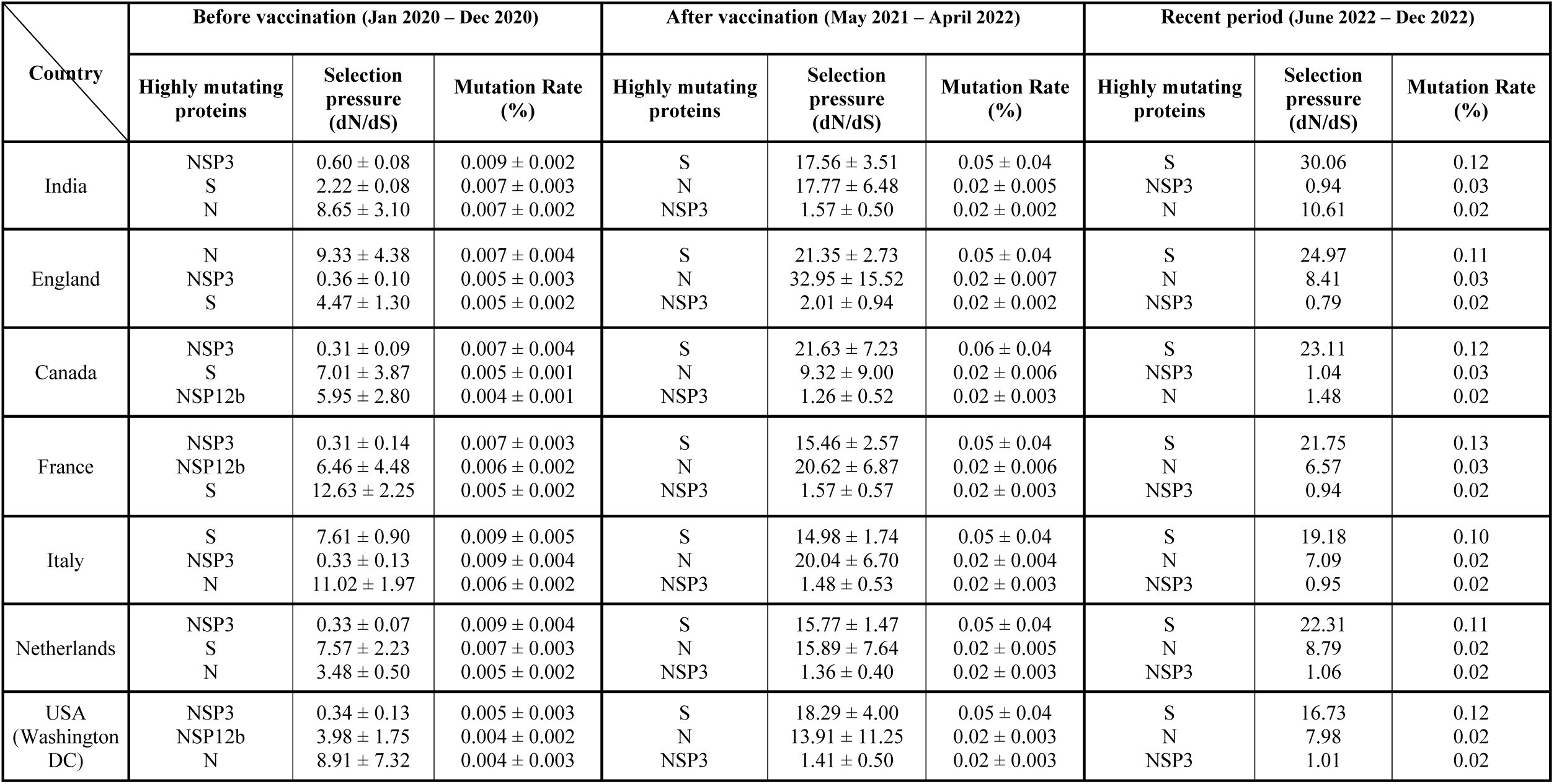
Data comparison of major mutating genes before, after vaccinations and recent period in each country

In RNA viruses, proofreading incapability of RNA polymerases accounts for their high mutation rates. Natural selection further results in increased adaptability and faster replication of viruses referring to their high mutation rates (40, 41). However, the highly conserved nsp14 in the *Coronaviridae family* has a proofreading function that may be a crucial factor for the large and complex viral genome (42). Absolute estimates of mutation rates in coronaviruses ranged from 0.67-1.33 × 10^-5^ per site per year in infectious bronchitis virus to 0.44-2.77×10^-2^ per site per year in mouse hepatitis virus. Comparatively, the global SARS-CoV-2 mutation rate was estimated to be moderate at 6 × 10^−4^ per site per year (43). The more than two-fold increase in mutation rates with time in SARS-CoV-2 as observed in this study would bring the genomic mutation rates at par with the reported genomic mutation rates for non-coronaviruses and the other RNA viruses like Influenza virus and HIV-1 (44).

### Selection pressure on the SARS-CoV-2 genome increased globally with time

The dN/dS ratio in each country for the pre-vaccination, post-vaccination and the recent period are listed in Table 1. The global dN/dS ratio in the SARS-CoV-2 genome was calculated as 1.72 ± 0.28, thus indicating a positive selection pressure. Our results are in line with a previous study that estimated the SARS-CoV-2 genome samples within the period of March-June 2020 to be under positive selection with a dN/dS value of 1.47 (45). The dN/dS ratio increased to 3.53 ± 0.98 in the post-vaccination period in various demographic regions. The recent period sequences showed an average dN/dS of 3.17 ± 0.15.

In Figure 1(B), comparison of dN/dS ratios in the countries for pre-vaccination, post-vaccination and recent period can be studied. There was more variation in the sequences taken from the post-vaccination period in comparison with dN/dS values before vaccination. Taken together, a definite increase in dN/dS is seen is post-vaccination sequences but the data from recent sequences was inconclusive because of small sample size and large standard deviation. The differences in the mean dN/dS ratios were found to be extremely statistically significant between the three phases with a two-tailed with p-value of less than 0.0001 (Table 2).

Previous studies reported that vaccinated individuals had significantly higher dN/dS than unvaccinated individuals and was calculated as 3.41 (46), which is similar to the calculated values in this study for post-vaccination and recent period. The increase in dN/dS depicts strong positive selection favouring dN in the SARS-CoV-2 genome (47, 48). In a previous study, it has been suggested that increase in the dN/dS ratios can cause increase in the mutation rates measured (49), which is also justified in our study as both mutation rates and dN/dS ratios are increasing after vaccination. Also, vaccination against SARS-CoV-2 in mass population is likely to have increased the selection pressure. Combined with the persistence of recurrent infections in immunocompromised individuals, this may have induced the selection of viruses with lower pathogenicity or virulence and higher transmission (50).

### Decrease in Ti/Tv was observed in SARS-CoV-2 genomes was observed with time

The estimated Ti/Tv ratios for each country in the pre-vaccination, post-vaccination, and the recent period are listed in Table 1. The Ti/Tv ratio in the pre-vaccination dataset was found to be 3.77 ± 1.10. The value is in agreement with the range of Ti/Tv of 2.0 to 5.5 calculated in previous studies (51, 52). The Ti/Tv ratio decreased in the post-vaccination period to 2.00 ± 0.12 and remained similar with the recent period sequences, i.e., 2.15 ± 0.05. The differences in the mean Ti/Tv ratios were found to be highly statistically significant between the three phases with p-value of less than 0.0001 (Table 2).

Figure 1(C) shows the comparison of Ti/Tv ratios within the countries in pre-vaccination, post-vaccination, and recent period. As observed from the figure, the proportion of Tv were found to be increased post-vaccination. As Tv are said to have a dominant contribution towards dN within the genome, an increase in Tv can be directly related to increase in dN. Thus, a decrease in Ti/Tv ratio is correlated with an increase in the dN/dS ratio. Previous studies have shown that Ti saturate more rapidly with time in comparison with Tv, thus causing Ti/Tv ratios to decline with evolutionary time (53–56).

### Change in Ti and Tv patterns work to equalize the CpG ratios in SARS-CoV-2 genome

The percentage of Ti and Tv in the pre-vaccination, post-vaccination and recent period are shown in Figure 2(A), (B) and (C) respectively. The most frequent Ti and Tv observed was C→U and G→U respectively in the pre- and post-vaccination period. The C→U and G→U substitutions result in the deterioration of CpG content. In the post-vaccination period, the C→U substitutions decreased by 12% from the pre-vaccination period, whereas in recent period, a slight decrease of 4% was observed from the post-vaccination period. In comparison with C→U, G→U substitutions deteriorated at a slower rate, i.e., 5% reduction post-vaccination and about 7% reduction in recent period from post-vaccination period. In the recent period sequences, C→U remained the most frequent Ti whereas U→G was the most frequent Tv.

**Figure 2:**
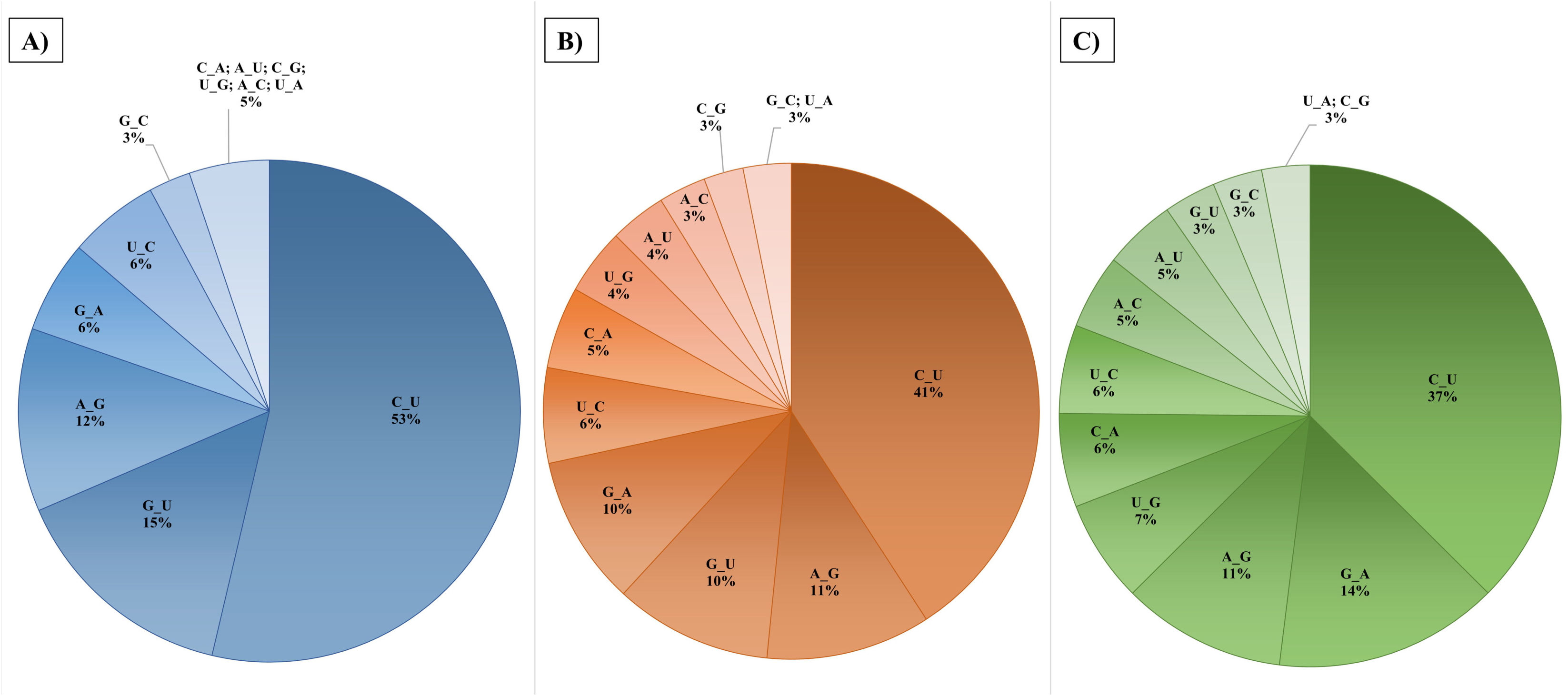
Percentage proportions of transition and transversion mutations in the countries in the three phases. **A)** Before vaccination (pre-vaccination period); **B)** After vaccination (post-vaccination period); **C)** Recent period.

A previous study that evaluated the sequences of the period January 2020 to March 2021 reported a drastic increase in C→U and G→U substitutions in the initial phase of SARS-CoV-2 infection, which eventually stabilized at a point of time and then decreased till March 2021 (57). Table 1 shows the %GC content in the genome sequences from each country for pre-vaccination, post vaccination and recent period. It can be observed that the %GC content slightly reduced after vaccination, whereas negligible reduction was seen in the recent period genome sequences. As a result, the overall decline in GC content is minimal and appears to be stabilizing in future. This is confirmed by another study which depicted minute variations in the CpG content in the SARS-CoV-2 genome sequences and concluded that CpG content reduced at a faster rate in the initial time of evolution which may slow down to become steady with the evolution in human host (58). Thus, these mutations were attributed to rapid evolution after transmission to the human host (59–61). The differences in the mean % GC content were found to be highly significant as tested a two-tailed p-value of less than 0.0001 (Table 2).

It has also been reported that the CpG motifs are lost in the SARS-CoV-2 sequences, which may enable antiviral response escape through TLR7 (58, 62). Higher U→G has been reported in the SARS-CoV-2 genome sequences collected between January 2022 to July 2022 (63). This further suggested that the amount of C→U and G→U substitutions are likely to decrease upon increase in divergence indicating that some portion of these mutations are knocked out by purifying selection, irrespective of the origin of SARS-CoV-2 (57, 64). Reduction in C→U and G→U substitutions together with the increase in U→G may eventually lead to an increase in the CpG content in the SARS-CoV-2 genome. Higher CpG content has been linked to the attenuation of the virus (65, 66).

### Rise of % mutations rates in S, N, NSP3 and NSP4 in the post-vaccination sequences

The pattern of mutations in proteins within the SARS-CoV-2 genome sequences was analysed for each protein in the pre-vaccination, post-vaccination, and recent period (Table 3). As shown, NSP3, S, N and NSP12b coding regions had the highest % mutation rates in the pre-vaccination period. There was little variation observed in the highest mutating genes in different countries studied. Previously, NSP3 was reported as the mutation hotspot of SARS-CoV-2 genome and mostly found to be co-mutating with NSP12b due to its involvement in replication and transcription complexes (67–71).

**Table 3:**
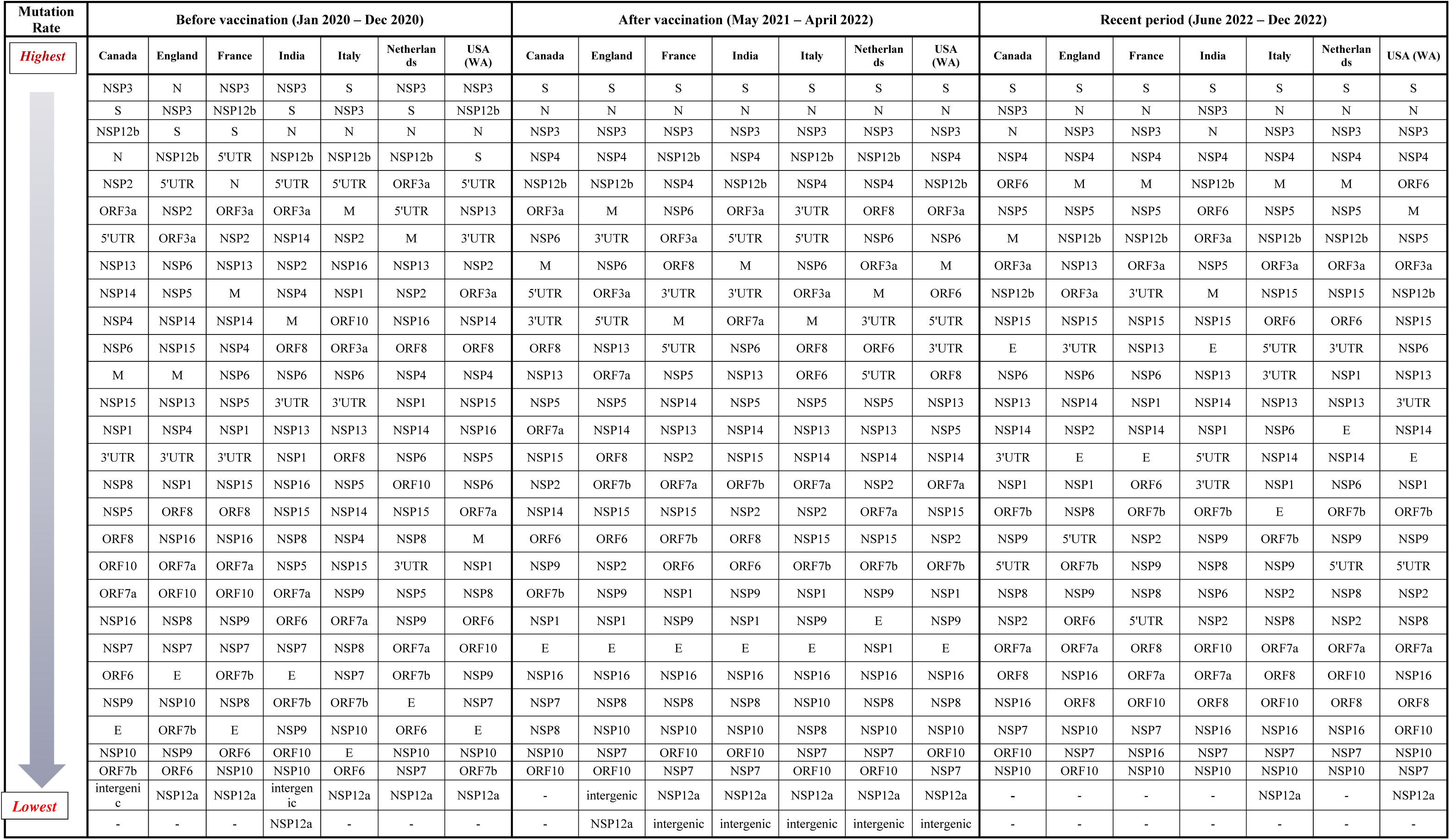
Protein-wise contribution to mutation rates in all the countries before, after vaccination and in the recent period

However, in the post-vaccination and recent period, the highest mutating genes remained the same, where S was observed to uniformly have the highest % mutation rates followed by N and NSP3. From the pre-vaccination sequences, S showed gradual increase in the % mutation rates till the recent period across all the countries. S contains the receptor-binding domain (RBD) which is highly variable, and through its mutational changes, it is known to affect viral replication, transmission and is also involved in immune escape (72, 73). In comparison with S, N has lower mutation rates and is comparatively stable (74). N is considered as a vital hotspot for mutations mainly in its serine-rich domain due to its involvement in viral replication and packaging (75, 76) and can be considered as a novel target for vaccine design (77).

Comparison of mutation patterns of all the proteins in Table 4 showed that NSP4 had shown low mutation rates before vaccination but rose to be the fifth highest mutating protein post-vaccination. In recent sequences, it uniformly occupied the fourth position and overtook NSP12b in all the countries. NSP4 along with NSP6 and NSP3 transmembrane proteins, induce the development of double membrane vesicles (DMVs) by reorganizing the endoplasmic reticulum of the host cell, and therefore it may be co-mutating with NSP3 (78, 79). In contrast with this, ORF10, NSP7 and NSP10 were the lowest mutating in all the three periods for all the countries taken into consideration. This was in line with previous studies where they were found to be conserved within the SARS-CoV-2 genome with few or no mutations (69, 80–82).

**Table 4:**
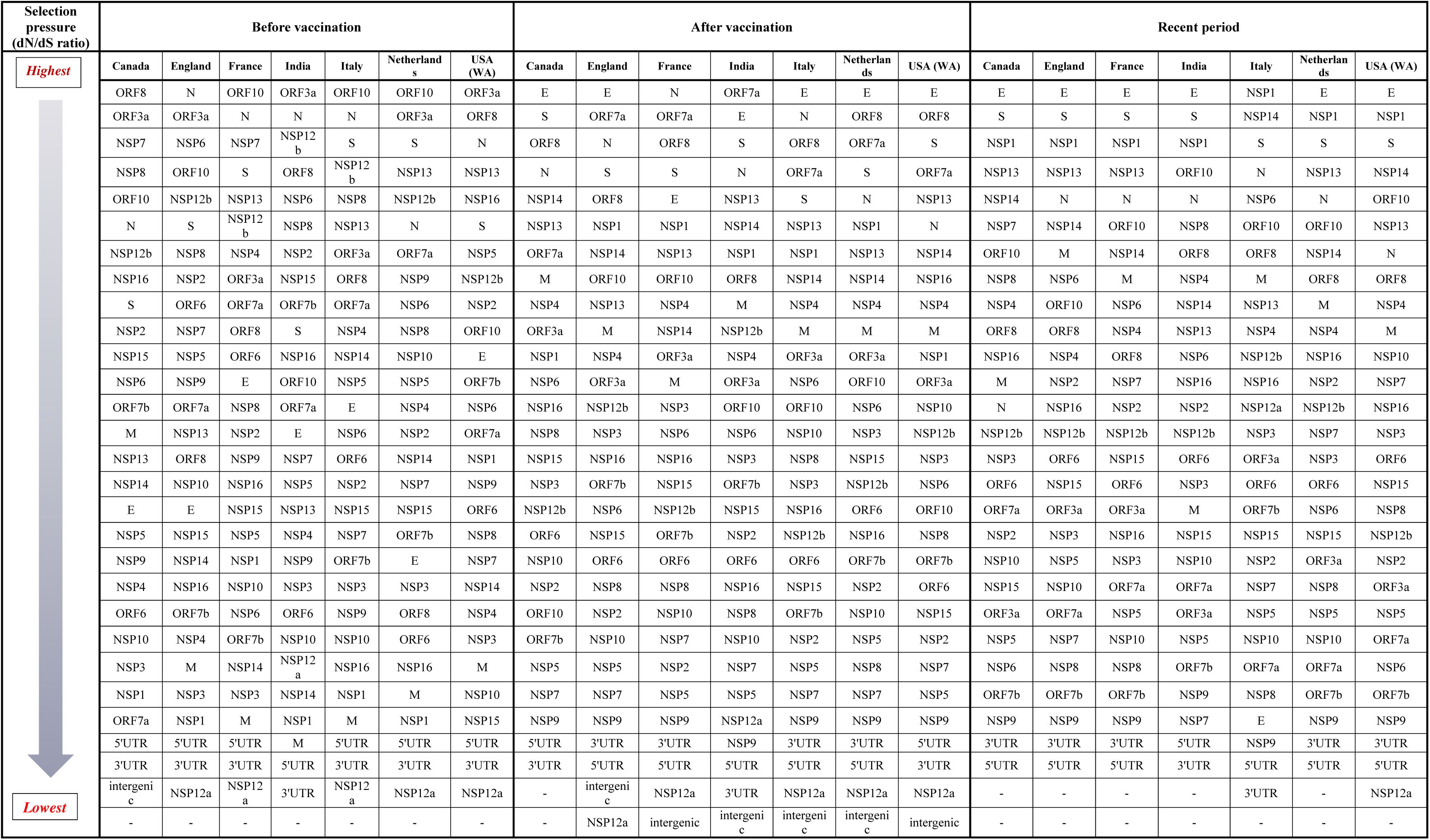
Protein-wise contribution to selection pressure (dN/dS ratio) in all the countries before, after vaccination and in the recent period

### Increase in selection pressure on E, S and NSP1 post-vaccination

The dN/dS ratios of the proteins in all the countries are shown in Table 5. SARS-CoV-2 proteins with the strongest positive selection pressures in the pre-vaccination period were N, ORF8, ORF3a and ORF10. In a change from the pre-vaccination period, the post-vaccination and recent phase showed E, S and NSP1 to be under high positive selection pressure with dN/dS much greater than 1.

**Table 5:**
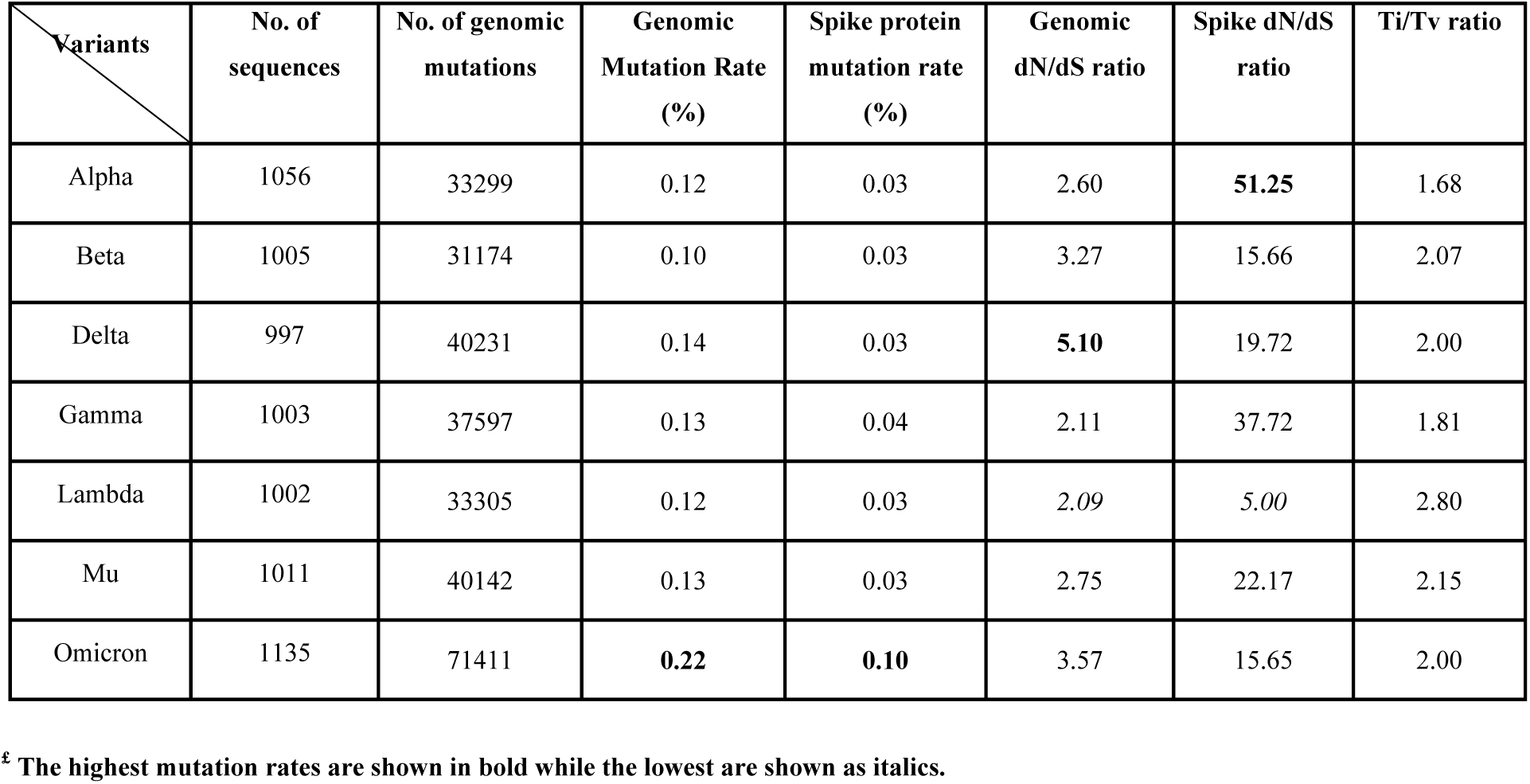
Comparison between genomic and dN/dS ratios in different SARS-CoV-2 variants

Previous studies during the year 2020 have identified ORF10 to be under strong positive selection, along with ORF3a, ORF8 and N also depicting much higher number of dN (83–85). ORF8 is involved in host-pathogen interactions through its 9 encoded proteins (86), ORF3a is involved in interference with host ion channels by encoding viroporin (87), while overexpression of ORF10 is known to downregulate IFN-1 expression, leading to suppression of the antiviral innate immune response (88). However, most viral proteins are multifunctional, and their other roles are likely to be discovered in future. In the pre-vaccination phase, NSP1 was under negative selection pressure with dN/dS<1. This was also shown in a previous study where NSP1 showed negatively selected sites only and it is designed to be evolutionary conserved for its functional requirement of host ribosomal complex binding (89, 90).

E and S indicated strong positive selection pressures in a 2021 study (91). E is involved in the expression of a small multi-functional protein that has an important role in host and virus interaction through ion-channel activity. Mutations within E are associated with a reduction in virulence (92, 93). S has shown continuous strong positive selection in our study in all the three phases as supported by other studies (48, 94).

NSP1 showed a large jump in the dN/dS ratio with a drastic increase in the number of dN after vaccination. NSP1 is suggested to suppress the host innate immunity by interacting with the 40S ribosomal subunit (90, 95, 96). NSP9 was observed to be the most common gene having dN/dS<1 showing purifying selection. NSP9 is said to be highly conserved and involved in viral RNA synthesis (97).

### Omicron genome had largest number of mutations, but dN/dS was highest in Delta

The genomic mutation rates (%), dN/dS and Ti/Tv ratios were also determined for each SARS-CoV-2 variant (shown in Table 6).

a. <u>Mutation rate (%) analysis</u>: The variant-wise genomic mutation rate analysis data suggests Omicron variant has the highest mutation rates amongst all variants (0.22%) followed by Delta (0.14%). Beta variant displayed the lowest mutation rate (0.10%). Previous studies showed Omicron to have high genomic mutation rate along with the accumulation of highest number of mutations (98–100).
b. <u>dN/dS ratio analysis</u>: As dN/dS ratios exceed unity for each SARS-CoV-2 variant, positive selection is responsible for promoting dN in the sequences. From Table 5, the Delta variant had the highest dN/dS ratio (5.1), followed by Omicron variant (3.57). Although the number of mutations is found to be higher in Omicron variant, the ratio of dN/dS is higher in the Delta variant. Delta variant was found to be highly virulent in comparison to the Omicron variant, which evolved to be more transmissible and less virulent (101–104). Therefore, the genomic dN/dS seems to correlate well with increased virulence. It has been argued that as virulence causes hindrance of transmission between hosts, dN/dS ratios may work to decrease virulence - thereby increasing transmission between hosts (105, 106).
c. Virulent genes have been observed to be highly subject to dN and therefore are under strong positive selection (107). Therefore, the protein-wise contribution to dN/dS was also determined in each of the variants and shown in Table 7. The unique genes having highest dN/dS in the highly virulent Delta variant were ORF7b and ORF7a while the unique genes contributing most to dN/dS in the Omicron variant were E, NSP14 and NSP1. The accessory protein ORF7b has been reported to mediate the cellular apoptosis caused by Tumour Necrosis Factor α.(108) The ORF7a protein initiates autophagy and helps in virus replication (109). Taken together, these proteins impair the host cell immune response and cellular function (110).
d. <u>Ti/Tv ratio analysis</u>: Lambda variant was found to have the lowest dN/dS ratio but the highest Ti/Tv ratio. A negative correlation has previously been observed between Ti/Tv ratio and dN/dS ratio under positive selection in the comparative analysis of SARS-CoV, SARS-CoV-2 and MERS-CoV for all the nucleotide substitution models (25). The reason for this could be explained by the fact that Tv substitutions favour dN. Previous studies have also remarked that higher number of Ti in the Lambda variant would remark lesser dN (53–56).

### S protein dN/dS ratios do not correlate with genomic dN/dS ratios in different variants

It has been suggested that the S mutations are positively selected to generate new variants of SARS-CoV-2 that have improved overall fitness (50, 111). To evaluate this, the average number of mutations, mutation rates and dN/dS ratios were determined for the S region in each variant. The average number of S mutations were calculated as a percentage of total number of S mutations in the genome. The resulting S mutations percentage for Alpha, Beta, Delta, Gamma, Lambda, Mu, and Omicron were 29.4%, 30.5%, 22.7%, 33.7%, 25.3%, 26.3% and 47.8% respectively. As seen in Table 5, the highest mutation rates in the whole genome as well as in S are seen in Omicron variant. S mutation rates of all the variants are similar except for Omicron, which has the highest S mutation rate of 0.1. This has also been observed in a previous study (98).

The dN/dS ratio of S was greater than 1 for all the variants. Alpha variant S protein showed the highest dN/dS ratio whereas Lambda variant S protein had the lowest dN/dS ratio. However, dN/dS values of both genome and S are comparatively lower in Omicron variant. Thus, there seems to be no correlation between dN/dS ratios of S and genomic dN/dS ratios, virulence, or transmission. That Omicron and Delta variant have the highest and the lowest S mutations respectively was also remarked in a previous study (112). The contribution of the SARS-CoV-2 proteins towards dN/dS ratios and mutation rates in each variant has been shown in Table 7 and 8 respectively.

## Conclusion

To the best of our knowledge, this is the only study that has compared the mutation rates, dN/dS ratios and Ti/Tv ratios during pre-, post-vaccination and the recent periods in different geographical areas after the emergence of COVID-19. While individual studies find support from literature, this remains a comprehensive study on genomic parameters of SARS-CoV-2 in different regions and periods.

The mutation rates were observed to be increased from the before vaccination period in each country to the recent period. In the pre-vaccination stage, the dN/dS ratio was greater than unity, showing mutations to be occurring because of natural selection. In the post-vaccination, it increased, signifying stronger positive selection in this period. However, the Ti/Tv ratio depicted significant decrease over time. Based on these parameters, the unpaired t-test helped in confirming that differences amongst the three phases were found to be extremely statistically significant (p-value < 0.0001).

The number of mutations has increased in S in the Omicron variant and accounts for 47.8% mutations in the genome. This might be a response to the vaccination as S is the main antigen target for the vaccine. However, most of them are dS mutations.

A higher dN/dS ratio is also seen in another structural protein N. A 2020 communication proposed N as a vaccine target in view of its low mutation rates (113). Indeed, the current diagnostic kits also target N recognition. This may need to be re-evaluated in the light of the high mutation rates as well as dN/dS ratios of N as observed in our study. Conversely, ORF10, NSP7 and NSP10 coding genes had the lowest mutation rates across all the considered demographic regions and could further be targeted and explored for SARS-CoV-2 diagnostics and therapeutics.

Amongst the SARS-CoV-2 variants, Delta and Omicron had the highest dN/dS ratio and mutation rates respectively. The Delta variant has dN and correlates with high virulence in comparison with the Omicron variant, which is less virulent but has higher transmissibility. It was also observed that the pattern of highest to lowest gene-wise dN/dS ratios unique in each variant genome.

Lastly, it is remarked that the mutation of the virus is thought to be happening in tandem with the genetic makeup of the host. Yet, it was observed that the worldwide patterns of genome mutations in SARS-CoV-2 remain the same all over the world despite the different vaccine technologies used. It seems possible that the viral genomic level changes are influenced by the presence of other circulating strains of SARS-CoV-2 as well as other viruses present in the environment.

## Data Availability

Raw data were generated from GISAID. Derived data supporting the findings of this study are available from the corresponding author SB on request.

The findings of this study for pre-vaccination period are based on metadata associated with 34970 sequences available on GISAID up to December 31, 2020, and accessible at doi:**10.55876/gis8.230517nt**

The findings of this study for post-vaccination period are based on metadata associated with 36020 sequences available on GISAID from May 15, 2021, to June 15, 2022, and accessible at doi:**10.55876/gis8.230517va**

The findings of this study for recent period are based on metadata associated with 3880 sequences available on GISAID from June 15 2022 to December 5, 2022, and accessible at doi:**10.55876/gis8.230518oq**

The findings of this study for pre-vaccination period are based on metadata associated with 7209 sequences available on GISAID from December 30, 2019, to January 9, 2022, and accessible at doi:**10.55876/gis8.230518xv**

## Acknowledgements

We gratefully acknowledge all data contributors, i.e., the Authors and their Originating laboratories responsible for obtaining the specimens, and their Submitting laboratories for generating the genetic sequence and metadata and sharing via the GISAID Initiative, on which this research is based.

## Abbreviations

SARS-CoV-2: Severe Acute Respiratory Syndrome Coronavirus 2
ACE2: Angiotensin Converting Enzyme 2
GISAID: Global Initiative on Sharing All Influenza Data
NSP: Non-structural protein
ORF: open reading frame
S: Spike protein
E: Envelope protein
M: Membrane protein
N: Nucleocapsid protein
dN: non-synonymous mutations
dS: Synonymous mutations
Ti: Transitions
Tv: Transversions
Ti/Tv: Transition to transversion ratio

